# Predicting Protein Binding Affinity With Word Embeddings and Recurrent Neural Networks

**DOI:** 10.1101/128223

**Authors:** Carlo Mazzaferro

## Abstract

At the core of our immunological system lies a group of proteins named Major Histocompatibility Complex (MHC), to which epitopes (also proteins sometimes named antigenic determinants), bind to eliciting a response. These responses are extremely varied and of widely different nature. For instance, Killer and Helper T cells are responsible for, respectively, counteracting viral pathogens and tumorous cells. Many other types exist, but their underlying structure can be very similar due to the fact that they all are proteins and bind to the MHC receptor in a similar fashion. With this framework in mind, being able to predict with precision the structure of a protein that will elicit a specific response in the human body represents a novel computational approach to drug discovery. Although many machine learning approaches have been used, no attempt to solve this problem using Recurrent Neural Networks (RNNs) exist. We extend the current efforts in the field by applying a variety of network architectures based on RNNs and word embeddings (WE). The code is freely available and under current development at https://github.com/carlomazzaferro/mhcPreds

## Introduction

The modeling of protein-protein interactions has seen recently a tremendous influx of computational approaches mainly due to its reduced cost as compared to *in vitro* testing [8] [4]. The ability to quickly model an interaction and its effect on the human body without running an entire experiment will result in reduced human labor as well as saved time for the results to come out. This is currently an unmet need in the field and could have long-lasting effects in this field of research. These methods could not have come without the increased data availability that has marked the past few decades and the increased computational power marked by a concurrent decrease in its cost. A remarkable effort has been made by organization such as IEDB [13] to curate and distribute data sets containing high-quality samples. The trend has been of continuous increase in data quality and size, which further solidifies the potential for the application of computational approaches to this field. In this paper, we introduce a command-line based python API for generating such predictions with accuracy comparable to state-of-the-art methods. At the time of this writing and to the best of our knowledge, no published peer-reviewed article exists describing an open source python framework for such task. The existing frameworks, especially the most widely used and accepted, are based on REST APIs or standalone programs where the source code is not accessible to the general public (see, for instance, http://www.cbs.dtu.dk/services/NetMHCcons/) [7]. The intent of providing a transparent and modifiable API serves the purpose of prompting others to engage in the effort of producing better algorithms and enabling collaborative efforts with standardized, cutting edge data analysis methods.

## Problem Overview

The main function of MHC molecules is to bind to antigens derived from pathogens and display them on the cell surface for recognition by the appropriate T-cells. The availability of the sequence data of HLA-binding peptides in the early 1990s [10] led to a search for commonalities among these sequences — that is, allele-specific motifs that convey binding. It quickly became clear that the interaction between HLA and peptides is rather complex, and thus more involved pattern-recognition methods were developed. The data that this paper is concerned consists of a list of peptide strings with a IC50 (i.e., the half maximal inhibitory concentration, a measure of the effectiveness of a substance in inhibiting a specific biological or biochemical function). These values range from a maximum of 50,000 nM (indicating low binding affinity) to close to zero, indicating very high binding affinity (minimum value is 0.788 nM). A successful predictive model to be applied to the data set in question should be able to regress, given a sample of peptide-IC50 scores, to an accurate prediction of the resulting IC50 score for novel peptide sequences.

### Past and Current Efforts

Supervised learning methods have been applied to this problem extensively. Initial efforts include position-specific scoring matrices (PSSM) [4], which represent a tool for modeling the probability of a sequence being biologically active, Support Vector Machines, Hidden Marakov Models, and more recently artificial neural networks (ANN) [10]. The evolution in the methods has seen a shift from linear models (PSSM) to highly non-linear ones such as neural nets. This trend is typical of many other fields that have seen an increase in data availability and compute power to develop more complex models. In this paper, we focus on extending the results obtained by applying ANN to this field by implementing a novel neural network architecture that learns deeper relationships between the positional information of amino acids (AAs) in a peptide sequence.

### Recurrent Networks

RNNs [11] are a family of neural networks for processing sequential data. Much as a convolutional networks is a neural network that is specialized for processing a grid of values such as an image, a recurrent neural network is a neural network that is specialized for processing a sequence of values *x*_1_ …, *x*_*τ*_. A key factor that determines the effectiveness of RNNs when dealing with sequences is parameter sharing [12]. A traditional fully connected feed-forward network would have separate parameters for each input feature, so it would need to learn all of the rules of the language separately at each position in the sentence. By comparison, a recurrent neural network shares the same weights across several time steps [3]. A particular subclass of RNNs are the so called Long-Short-Term Memory (LSTM) [5] networks. Unlike traditional RNNs, an LSTM network is well-suited to learn from experience to classify, process and predict time series when there are time lags of unknown size and bound between important events. Relative insensitivity to gap length gives an advantage to LSTM over alternative RNNs and other sequence learning methods in numerous applications. In this paper, we explore the effectiveness of LSTMs in identifying higher order interactions between amino acids and their positional information within a peptide string.

### Embedding Networks

Another approach that was be taken is the development of a network architecture based on WE. Simply put, this set of techniques are generally characterized by the mapping from a set of words or phrases from the vocabulary to vectors of real numbers. The reason for including this approach is the interest in contrasting and comparing different, although related methods. In particular, RNNs main strength is the ability to learn representations from sequentially related inputs. Word embeddings, on the other hand, code these relationship in the data itself. More specifically, using word embeddings results in the extraction a meaningful representation from a sequence of letters by finding a transformation from a sparse representation to a a denser one. In particular, word embeddings aim at coding for more meaning positional information about a sequence. Word2Vec [9] represents one of such efforts applied to natural language processing (NLP). A particularly key insight drawn from the this paper is that the vocabulary by which our language (which can mapped to the set of peptides to be analyzed, for our present problem) are encoded as discrete entities that appear sequentially. The key aspect of Word2Vec, however, is transforming such mapping into a continuous space. This allows to use continuous metric of similarity to evaluate the semantic quality of our embedding. The end goal is to, by using a continuous representation, mapping similar words (amino acids) to similar regions (peptides). This mapping is done through what in most deep learning APIs is called an embedding layer.

### The Data

Data was gathered from IEDB’s website [8]. It contains MHC-peptide binding affinity measures for a variety of different species and alleles. It must be noted that the affinities are very much dependent on the structure of the MHC itself, whose structure varies widely depending on the allele and the species in question. In particular, we are interested in the subset of human alleles for which the number or samples is large enough to build a robust model. The total number of samples in the data set is of 176,161, out of which most are from human alleles. However, since there are a grand total of 118 known alleles in the data set, building a robust model can be challenging due to the limited samples per allele. On average, for the human samples, there are a total 900 samples. We selected the alleles for which most data samples are collected. A set of 6 alleles (HLA-A0201, HLA-A0101, HLA-A0301, HLA-A0203, HLA-A1101, HLA-A0206) was used to train the models and determine optimal parameters, although any allele can be trained and predicted on. In order to extract maximal information from the peptide sequences, two approaches were taken: ‘one hot encoding’ each peptide as a *x* by 20 binary matrix, where x is the length of the peptide, and 20 is the number of possible AAs. Using this method, the matrix will contain as many ones as AAs, where each one denotes the positional information of the AA within the peptide string. For example, the peptide “STAA” would be encoded as (the transpose of) the following matrix:

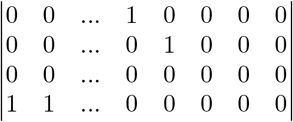

This representation however, is remarkable for its highly sparse nature. Out of the 180 entries, only four are non-zero. Despite the known difficulty of training deep learning models in data of this kind [2], this representation was chosen due to the interest in using RNNs and to test its effectiveness and robustness for the task of learning higher dimensional representations of peptide sequences, and in particular the positional interaction between them. Another encoding method used was the sparse k-mer encoding method, based on the previously described WE. This methods simply assigns an index value to each amino acid in a peptide according to a predefined dictionary of amino acids. For instance, the peptide “STAA” would be represented as:

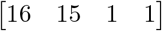

Another step taken was to normalize the input sequences to a single fixed length. For the purpose of this experiment, the peptide length was restricted to 9 AAs. Peptides of varying length were processed and extended/reduced in size by slicing them into sub strings of fixed length and applying either the index or one encoding to these sub strings. If a string was longer than the desired length, then it was reduced to the desired length by deleting characters at all possible positions.

## Methods

Two sets of experiments were performed. The first one was based on the application of kmer embedding on a simple two-layer neural network in order to create a performance baseline and validate the approach. The second experiment, which was based on the application of a variety of different RNN architectures, was design to extend and improve currently existing methods. Tensorflow [1] was used, alongside with a Tensorflow-based higher level API for rapid prototyping named tflearn. An AWS instance running Tensorflow 1.0 on a NVIDIA K520 GPU was used for most computations. Tensorboard was used to monitor progress at early stages to ensure learning was consistent through the epochs, and later used as well to generate the visualizations of the network such as the one seen in Figure 1. In both experiments, the IC50 targets were mapped to the range {0, 1}, so that the output from a sigmoid activation function would be meaningfully comparable to the targets of the problem.

**Figure 1.**
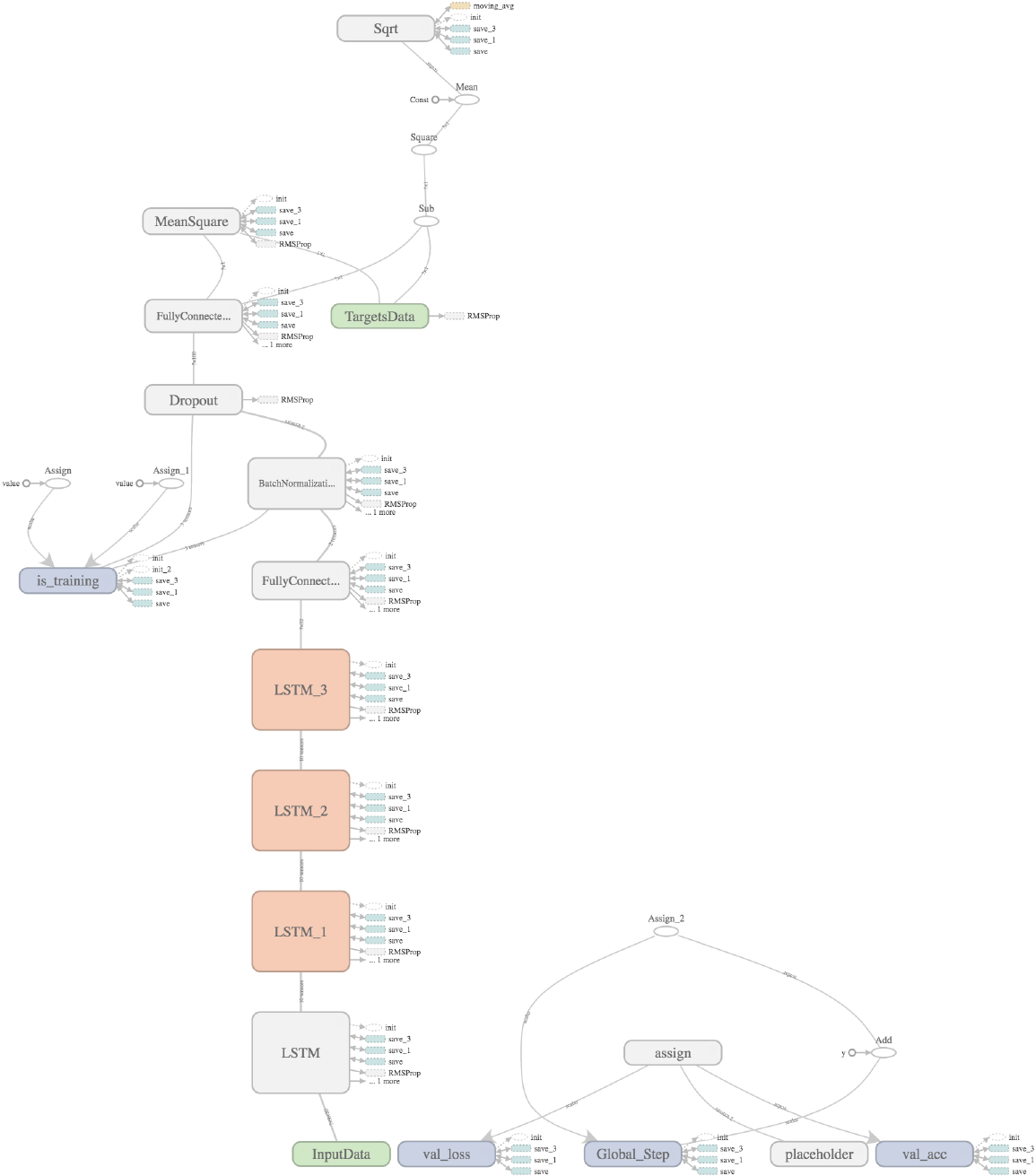
Deep LSTM Neural Network.

### Hyperparameter Search

Tuning parameters for a specific architecture may be a daunting task. The number of hyper parameters is exceedingly large, and in order to develop a computationally feasible approach, some limitations had to be taken. In particular, the optimizers used for the architectures, as well as the loss function, batch normalization rounds, and number of epochs were kept constant. This heuristic was determined as efficient solely after multiple rounds of training and evaluation performance. It was noted that these set of parameters lead to the most robust architectures, and were thus kept consistent through the experiment. Note that the general user can tune and modify them as desired through command-line inputs. For the remaining hyperparameters, a brute-force approach was taken. Namely, a set of 3-5 values for each of learning rate, batch sizes, and number of deep layers (for the RNN), embedding output size (for the WE network) were swept through iteratively and their accuracy calculated for each combination of hyper parameter. Incidentally, the size of the data sets was small enough to permit a thorough evaluation of each metric. A key point to be made is that the models were trained on subsets of the whole data set corresponding to each of the specific alleles previously mentioned, meaning that each hyperparameter search was performed a total of six times for each of networks. The results were written to a specific folder were a csv file with the predictions was stored alongside with a text file containing details about the run and the parameters used. A bash script was used to perform this steps without the need of the user’s supervision. A total of 360 runs was performed for the RNN, and 120 for the WE network. A variety of accessory scripts and classes were implemented to efficiently extract the meaningful data generated from these runs.

## 1 Results

Performance evaluation was done by plotting the calculating the area under the ROC curve [6] for the predictions. Adopting the same method as Nielsen et. al, [10], we determined the number of alleles having strong binding activity (IC50 below 500 nM) and calculated the number of true and positives false positives of our predictions for each class (either below or above 500 nM). Figure 2 shows the plots constructed from the predictions coming from each architecture on the six specified alleles using the optimal parameters found through the previously described brute-force search.

**Figure 2.**
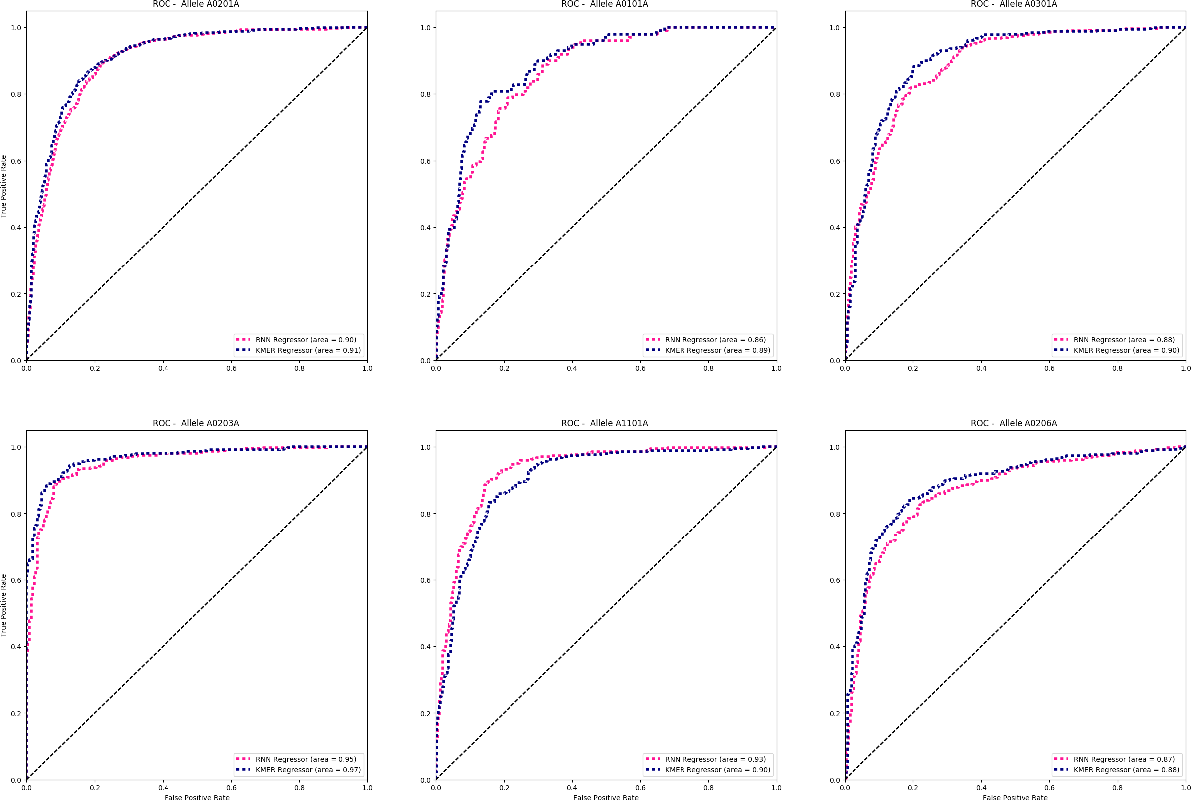
ROC Curves for the top predictors for from each network architecture on the six selected alleles.

Remarkably, both networks performed very similarly on the task. The network based on WE was initially set up as a standard neural network with two hidden layers and 100 hidden units, later increased to 150. The RNN instead achieved its best performance with a lower learning rate than the WE network and with two added deep LSTM layers. Despite the remarkable difference in how the input data was fed into the systems and how peptides were encoded into it, both architectures were able to learn much of the relationships between peptides and their positions exceedingly well. Something that should be noted, however, is the fact that the AUC values were heavily dependent on sample size. This result clearly shows that although a deeper net may be able to learn more complex representation given less data, sample size will always be amongst the most important performance-determining factors.

**Table 1.** Hyperparameters for nest run on allele A0203. AUCs were 0.9667 and 0.9543.

| Parameters | Best: RNN | Best: WE Net |
| --- | --- | --- |
| Learning Rate | 0.001 | 0.01 |
| Batch Size | 70 | 80 |
| Deep Layers | 2 | NA |
| Embedding Size | NA | 32 |

## 2 Discussion

We here presented a framework for the development of computational studies targeting novel drug-discovery using a novel approach to protein-protein representation, interaction, and modeling. It is designed to be extensible, hackable, and flexible enough to allow researchers to both use it a tool for generating new predictions from their own sequences, as well as deep learning enthusiasts interested in novel, unexplored applications of such techniques. State-of-art performance was achieved in a subset of the data using a novel architecture which still has plenty of room for improvement. Future efforts will be targeted towards integrating different representations of the data with architectures that leverage such representations. In particular, the inclusion of other biologically relevant data, such as protein 3D structure, as well as larger data sets to enable robust training, will indubitably mark the advancement of this field enabling cheaper, more specialized therapeutic and medical solutions.

## Acknowledgments

We thank Zhuowen Tu and Teaching Assistants for the UCSD COGS 181 course for the continuous support, feedback and knowledge shared; Kathleen Fisch and the Center for Computational and Bioinformatics at UCSD for the biologically relevant insights; and finally the AWSedu and the COGS department for letting us use their computation resources.

